# Systematic cross-species comparison of prefrontal cortex functional networks targeted via Transcranial Magnetic Stimulation

**DOI:** 10.1101/2023.12.20.572653

**Authors:** Taylor Berger, Ting Xu, Alexander Opitz

## Abstract

Transcranial Magnetic Stimulation (TMS) is a non-invasive brain stimulation method that safely modulates neural activity in vivo. Its precision in targeting specific brain networks makes TMS invaluable in diverse clinical applications. For example, TMS is used to treat depression by targeting prefrontal brain networks and their connection to other brain regions. However, despite its widespread use, the underlying neural mechanisms of TMS are not completely understood. Non-human primates (NHPs) offer an ideal model to study TMS mechanisms through invasive electrophysiological recordings. As such, bridging the gap between NHP experiments and human applications is imperative to ensure translational relevance. Here, we systematically compare the TMS-targeted functional networks in the prefrontal cortex in humans and NHPs. To conduct this comparison, we combine TMS electric field modeling in humans and macaques with resting-state functional magnetic resonance imaging (fMRI) data to compare the functional networks targeted via TMS across species. We identified distinct stimulation zones in macaque and human models, each exhibiting variations in the impacted networks (macaque: Frontoparietal Network, Somatomotor Network; human: Frontoparietal Network, Default Network). We identified differences in brain gyrification and functional organization across species as the underlying cause of found network differences. The TMS-network profiles we identified will allow researchers to establish consistency in network activation across species, aiding in the translational efforts to develop improved TMS functional network targeting approaches.

## Introduction

Transcranial magnetic stimulation (TMS) is a non-invasive brain stimulation method that can safely modulate neural activity in vivo (Rossi et al., 2009). This technique applies a rapidly changing magnetic field through a coil placed at the scalp, which induces an electric field in underlying brain regions. TMS has high spatial resolution (Deng et al., 2013; Opitz et al., 2011), which enables targeted stimulation of specific brain networks. As such, TMS is an emerging therapeutic option for several neurological and psychiatric disorders (Lefaucheur et al., 2014). TMS is approved by the Food and Drug Administration to treat depression, obsessive-compulsive disorders, and smoking cessation, with ongoing clinical trials for other indications. Despite its increasing applications in clinical and basic research, the physiological outcomes of TMS are known to be variable across individuals (Hamada et al., 2013; López-Alonso et al., 2014). One potential reason is the imprecise targeting of brain circuits with TMS. Due to individual differences in neuroanatomy, the induced electric fields can differ significantly across individuals. To address this issue, computational models based on the finite element method (FEM) have been developed to estimate the TMS-induced electric field in individually realistic head models (Windhoff et al., 2013). This modeling technique allows researchers to predict the impact of unique anatomical patterns, such as brain gyrification and cerebrospinal fluid (CSF) thickness, on the stimulation regions (Opitz et al., 2011; Thielscher et al., 2011). However, acknowledging that the correspondence between anatomy and function is not always one-to-one, FEM modeling, while adept at capturing unique anatomical features, does not incorporate aspects of functional brain organization. This is particularly important for targeting higher-order association areas implicated in the treatment of depression (Rizvi and Khan, 2019; Schutter, 2009; Zhao et al., 2022).

In higher-order association areas, the relationship between brain anatomical landmarks and functional behavior remains subject to ongoing research (Amiez et al., 2006; Goulas et al., 2012; Margulies and Petrides, 2013). Resting-state functional magnetic resonance imaging (r-fMRI) has emerged as a popular method to map functional brain networks in vivo (Power et al., 2011; Stephen M. Smith et al., 2013). R-fMRI enables precision mapping of individual functional networks with high test-retest reliability (Gratton et al., 2020; Laumann et al., 2015). Consequently, researchers have suggested using r-fMRI to guide TMS targeting (Fox et al., 2012; Oathes et al., 2021). Network-guided TMS has the potential to account for individually unique functional brain networks and target symptom-specific brain networks (Siddiqi et al., 2020). Our research team has further developed this method and combined it with FEM modeling (Opitz et al., 2016), identifying distinct r-fMRI networks stimulated by TMS-induced electric fields. This allows researchers to integrate both anatomical and functional aspects of TMS targeting. While these methods have provided important tools to predict TMS activation of functional networks, they need further experimental validation. In humans, mostly indirect measurements of TMS physiological effects are available, hampering efforts to elucidate TMS network mechanisms. Research in non-human primates (NHPs) has proven fruitful in exploring these assumptions and studying TMS mechanisms leveraging invasive physiological recordings.

Non-human primates are ideally suited to investigate TMS neural mechanisms (de Lima-Pardini et al., 2023; Hanlon et al., 2021). Compared to other mammals, NHPs have human-like cortical complexity and prefrontal cortex development (Lear et al., 2022). These features allow researchers to conduct translational studies using TMS of prefrontal brain regions, a common target area in treating depression. More importantly, invasive recordings in NHPs have allowed researchers to study the neural effects of TMS with precision not available in humans (Mueller et al., 2014; Perera et al., 2023; Romero et al., 2019). However, despite its unique opportunities, the translation between NHP and human TMS applications is not straightforward. The anatomical features (i.e., size, gyrification) (Alekseichuk et al., 2019) and functional organization (Xu et al., 2020) lack one-to-one homology to human models. Notably, the evolutionary expansion of the human neocortex results in a highly convoluted cortex, especially in higher-order regions (Donahue et al., 2018; Mars et al., 2018; Van Essen, 2004; Van Essen and Dierker, 2007). Salience, frontoparietal (FPN), default mode networks (DN), and their interactions exhibit greater variations in humans than those observed in NHPs (Ardesch et al., 2019; Donahue et al., 2018; van den Heuvel et al., 2023). This disparity highlights the need for a bidirectional translational pipeline that can map TMS functional network targeting between humans and NHPs.

Here, we develop an integrated cross-species framework that combines interspecies anatomical alignment with r-fMRI to map TMS functional networks between humans and NHPs. This is based on our recently developed cross-species functional alignment method (Xu et al., 2020) that enables a quantitative comparison of functional homology across humans and NHPs. We focus on the prefrontal cortex due to its intricate complexity across species and crucial role in TMS clinical applications. We systematically compare the functional networks targeted with TMS in the prefrontal cortex between macaques and humans. We highlight commonalities and differences in TMS functional networks across species and investigate factors leading to these observed differences.

## Methods

### Overview

To compare TMS-activated functional networks in humans and non-human primates, we used two publicly available datasets of functional MR images: the Human Connectome Project (HCP) s1200 release (Van Essen et al., 2012) and the Oxford (anesthetized) macaque samples from the open-source NHP consortium, PRIMatE-Data Exchange (PRIME-DE) (Milham et al., 2018). For each dataset, we developed individual FEM head models and simulated the TMS-induced electric fields for a set of locations in prefrontal brain regions (**Figure 1A, C**). The TMS electric field simulations were used as weighted seed regions for functional connectivity analyses (**Figure 1B, D)**. We categorized which functional networks are activated by TMS using the Yeo parcellation (Yeo et al., 2011). Finally, we compared the functional network activation pattern across species by utilizing the previously established macaque-human cortical transformation (Xu et al., 2020).

**Figure 1.**
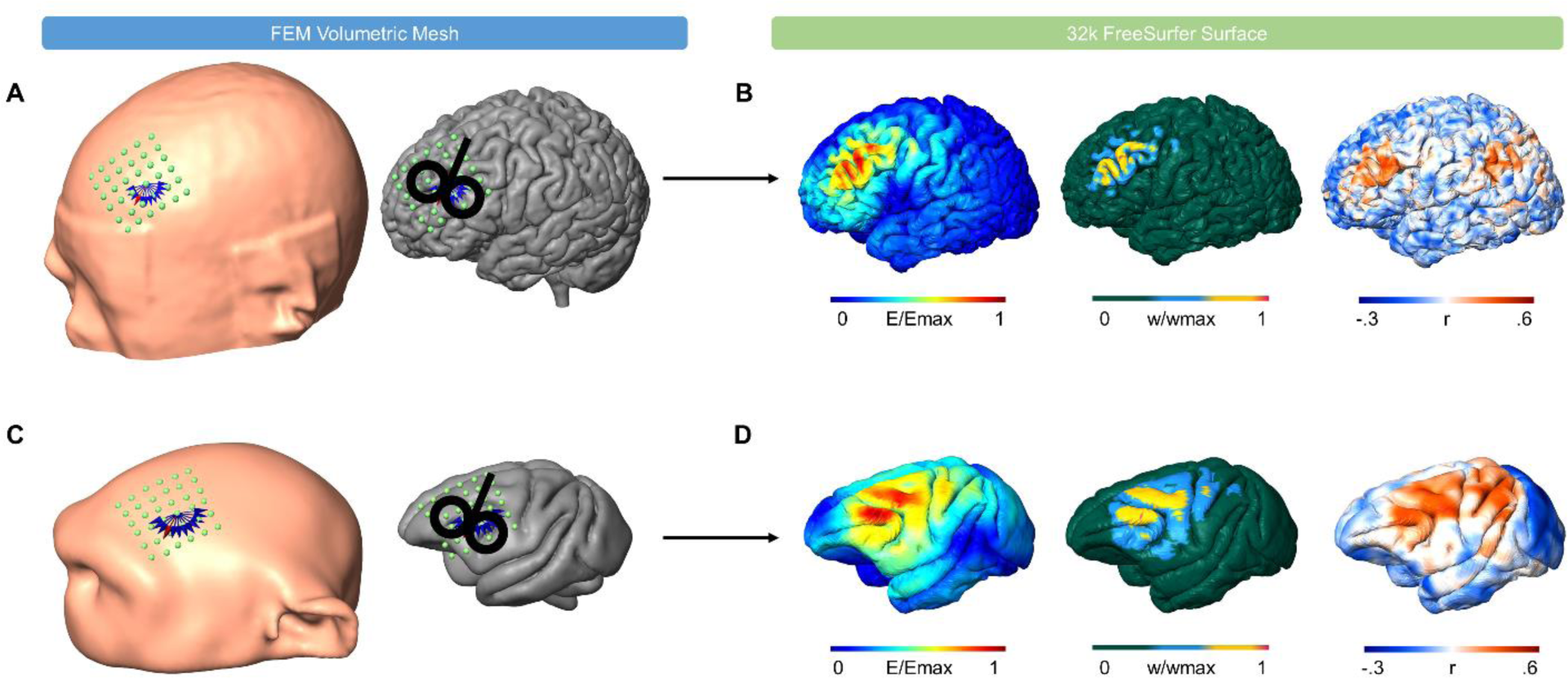
Overview of the integrated TMS-rfMRI network modeling pipeline. The TMS-fMRI network modeling pipeline predicts the ability of TMS to activate, at a specified coil location and orientation, specific functional connectivity networks. From anatomical MR images, finite element method (FEM) head models are generated for both human and non-human primate (NHP, macaque) datasets. **A)** Stimulation grids are placed over F3, depicted for one human subject. Each stimulation grid consists of 36 points (6×6, directions: lateral-medial, anterior-posterior with 10mm spacing) (left). For each grid location, 12 coil orientations (0-165 degrees, 15-degree steps) are simulated, shown at one sample location over the gray matter surface (right). **C)** Stimulation grids are placed over the left prefrontal cortex, consisting of 36 coil locations (6×6, directions: left-right, anterior-posterior) with 5mm spacing) 4 mm above the scalp, depicted on one NHP subject (left). For each coil location, 12 coil orientations (0-165 degrees, 15-degree steps) are simulated, represented in blue and red arrows at one sample location. The grid and the sample coil orientations are shown over the pial surface (right). **B, D)** The simulated electric field strength (E) is interpolated to the 32k FreeSurfer LR (fsLR) surfaces generated within the HCP preprocessing pipeline. An example electric field distribution (left) for the coil location and orientation (indicated by the red arrow in panel A) is depicted on the 32k fsLR surface. Node weights within the pial surface are created based on their electric field strength (threshold: > 50% Emax) to determine a seed region (middle). A functional connectivity map (right) for the seed region is generated by correlating at each pial node the r-fMRI time series with the weighted time series of the seed region.

### Human data

We selected MRI data from ten randomly selected, unrelated participants from the HCP s1200 release (Van Essen et al., 2012). Images included T1-weighted MP-RAGE and T2-weighted MP-SPACE MRI scans and a r-fMRI scan. The r-fMRI scan used within this analysis was acquired on day one of the HCP S1200 fMRI protocol (TR=0.72s, 2mm isotropic) and contained a single 15-minute run (phase encoding left-right) for each participant. Image acquisition and preprocessing protocols are available with the HCP S1200 release (WU_Minn, 2017). The Minimal Preprocessing Pipeline (MPP) (Glasser et al., 2013) was applied to all structural imaging files in the HCP preprocessing pipeline. Briefly, MPP includes spatial/artifact distortion removal, cross-modal registration, surface generation, and alignment to a symmetric fsLR-32k template surface (Van Essen et al., 2012). The functional MPP includes motion correction, distortion correction, ICA-FIX denoising, and spatial smoothness (FWHM=2mm) (Stephen M Smith et al., 2013). In our analysis, we used the 32k FreeSurfer LR (fsLR) aligned surfaces in native space for each participant, which capture the individual morphology of the brain shape. The aligned 32k fsLR surfaces contain 32,492 vertices per hemisphere that are comparable across participants.

### Macaque data

The rhesus macaque (*Macaca mulatta*) data were available from the Oxford dataset of the PRIME-DE consortium (Milham et al., 2018). The r-fMRI scan consisted of a single 53.3-minute run per animal under anesthesia with isoflurane. The ethics approval for animal care, MRI, and anesthesia was carried out in accordance with the UK Animals (Scientific Procedures) Act 1986. The details of the animal housing, anesthesia protocols, and MRI acquisition were reported in previous studies (Noonan et al., 2014) and PRIME-DE (Milham et al., 2018). The macaque imaging data underwent preprocessing using a customized HCP-like pipeline (Xu et al., 2019, 2018, 2015). Briefly, the preprocessing of r-fMRI includes slice timing, motion correction, co-registration, nuisance regression (Friston’s 24 motion parameters, mean signal of white matter (WM) and CSF), and band-pass filtering (0.01-0.1 Hz). Finally, the preprocessed data was projected from the volume to the surface space and smoothed (FWHM=2mm) along the surface. To ensure the accuracy of the individual FEM model, quality assessment and visual inspection were conducted on the co-registration and surface reconstruction steps. Ten animals were included in our final analysis. Like HCP’s MPP, 32k FreeSurfer surfaces were generated per hemisphere for each animal. The 32k fsLR surfaces enable direct comparison across animals within the macaque dataset.

### Estimation of TMS induced electric fields in Humans

We created individual FEM models for each of the ten participants based on the high-resolution T1- and T2-weighted images using SimNIBS 3.2 (Thielscher et al., 2015). We constructed a stimulation grid of 36 coil locations centered on F3 (6×6, left-right and anterior-posterior directions, 10mm spacing) on each FEM head model, as illustrated in **Figure 1A**. We designed the grid to cover different TMS targeting strategies for the dorsolateral prefrontal cortex (DLPFC) (Avery et al., 2006; George et al., 2010, 1995; Herbsman et al., 2009; Herwig et al., 2001; Pascual-Leone et al., 1996). We simulated 12 distinct coil orientations, stepping in 15-degree increments, covering a 180-degree half-circle. At 180-degree turns, electric fields are identical, with only the field direction reversed. We simulated TMS electric fields using a Figure-8 Magstim coil with a 70mm diameter. For each participant, we simulated 432 electric fields (36 coil locations x 12 coil orientations). We interpolated the calculated electric field strength from the FEM volumetric mesh to the pial surface of the 32k fsLR subject surface using an iterative closest point (ICP) algorithm. Briefly, the ICP algorithm calculates the transformation matrix between a three-dimensional (3D) source and a 3D target model (Rusinkiewicz and Levoy, 2001). In this study, we used the ICP-generated transformation matrix to perform the linear transformation between the gray matter surface of the FEM volumetric model and the pial surface of the 32k fsLR model. **Figure 1B (left)** depicts an example of a simulated electric field distribution.

### Estimation of TMS induced electric fields in Macaques

To create a realistic FEM head model, we manually segmented an anatomical T1 image from the Oxford dataset into six unique tissue types (skin, skull, CSF, GM, WM, Eye) using ITK-SNAP (Yushkevich et al., 2006). We used this segmentation to create a realistic volumetric head model using SimNIBS 3.2 (Thielscher et al., 2015). Due to the complexity of FEM head model generation in macaques and the requirement to manually segment each tissue layer, we constructed one complete macaque head model. We created a grid of 36 coil locations, organized in a 6×6 arrangement with directions spanning left-right and anterior-posterior, maintaining a 5mm spacing, illustrated in **Figure 1C**. The NHP grid was designed to mimic the grid used for the human model, scaled down to account for the smaller head size of the macaque. At each coil location, we investigated 12 unique coil orientations to cover a 180-degree half-circle with 15-degree incrementing steps. Using this configuration, we generated 432 unique coil parameter sets (36 coil locations x 12 coil orientations). We simulated TMS electric fields for the same TMS coil used for the human simulations (focal Figure-8 coil, 70mm diameter, Magstim). We interpolated the simulated field strength from the FEM volumetric mesh to the pial surface of the 32k fsLR surface of all subjects within the macaque dataset using an ICP algorithm. **Figure 1D (left)** shows an example of electric field distribution on the NHP 32k fsLR surface.

### Subject Level TMS Resting State Analysis

For each participant, we conducted 432 TMS electric field simulations and interpolated these results from the FEM volumetric head models to the 32k fsLR pial surface. For each simulation on the 32k fsLR pial surface, we determined an associated seed region by thresholding the electric field to greater than 50% of the maximum electric field strength of the pial surface (Opitz et al., 2016). Other electric field threshold values generate comparable results (**Supplementary Figure 1**). In the seed region, we assigned a weight to each node by normalizing the electric field strength at the node by dividing the node’s electric field strength by the total electric field within the seed area. Example seed regions for a human and macaque example case are shown in **Figure 1C** and **Figure 1D**. We computed a weighted-average time series by summing the fMRI time series at each node and multiplying by the individual node weight for each seed region (Opitz et al., 2016). We correlated the weighted-average time series with the time series of each node of the pial surface to calculate a whole-brain functional connectivity map. Example whole-brain functional connectivity maps are shown in **Figure 1C** and **Figure 1D** for both humans and macaques, respectively. To account for the effects of coil location, we calculated correlations using partial correlations, with the mean overall 432 averaged time series used as a covariate (Opitz et al., 2016).

### Parcellation of Functional Networks

We mapped the Yeo-7 functional network parcellations (Yeo et al., 2011) from FreeSurfer fsaverage surface to the HCP standard fsLR-32k cortical surface using the label-resample command (Glasser et al., 2013). We used this parcellation to identify different functional networks for each participant. The Yeo-7 parcellation mapped on an example 32k fsLR human surface is shown in **Figure 2A**. The corresponding Yeo parcellation map on the macaque 32k fsLR surface was generated via a previously established cross-species functional alignment (Xu et al., 2020). In short, this alignment was built based on the matched functional connectivity profiles between human and macaque monkeys, which quantifies homologous regions even when their location was decoupled from anatomical landmarks (Xu et al., 2020). The NHP Yeo-7 parcellation mapped on the 32k fsLR NHP surface is shown in **Figure 2C**.

**Figure 2.**
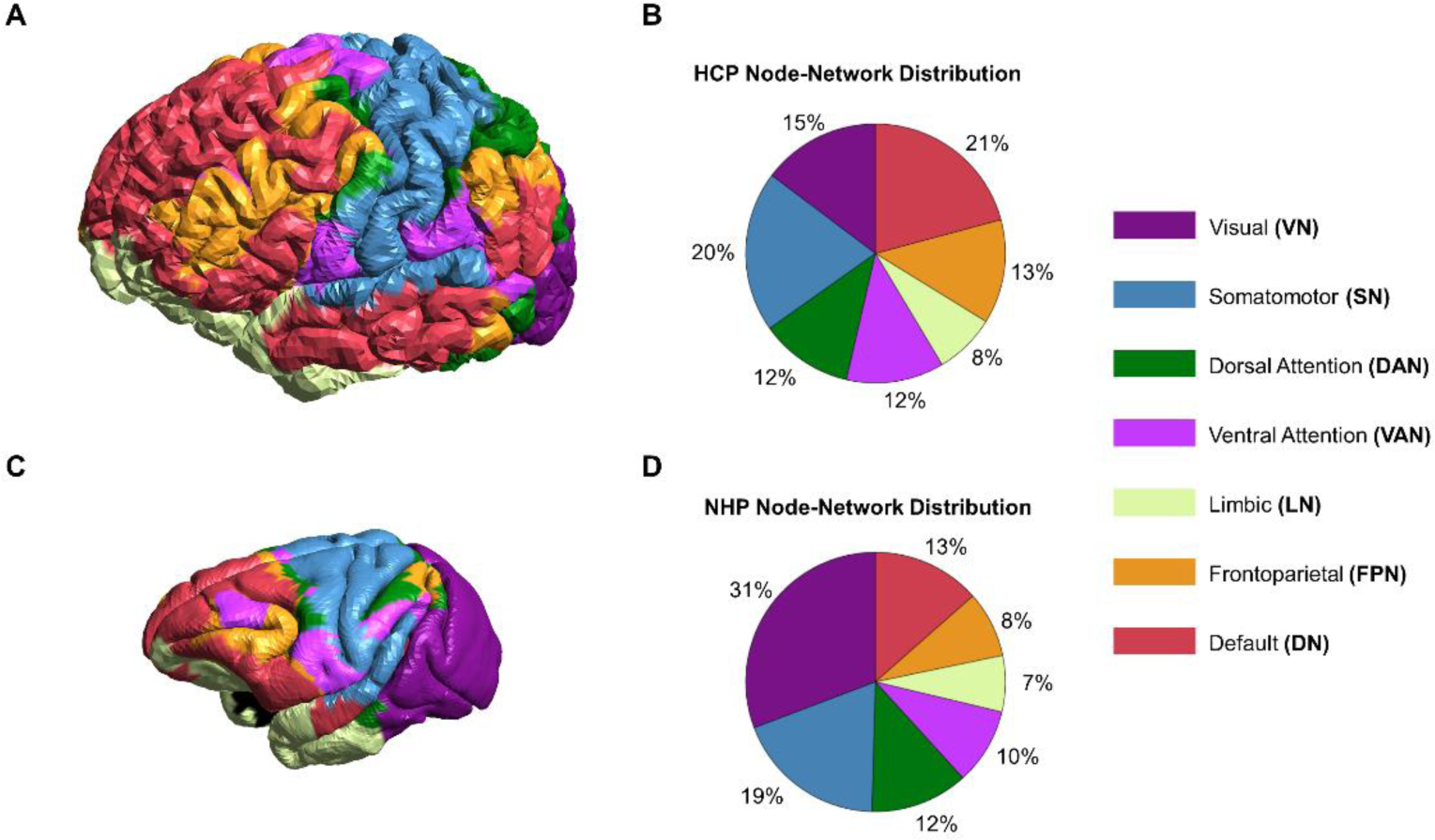
Yeo Functional Parcellation across species. The Yeo parcellation divides the gray matter surface into seven functional networks: Visual (dark purple), Somatomotor (blue), Dorsal Attention (green), Ventral Attention (light purple), Limbic (yellow), Frontoparietal (orange), Default (red). **A)** Yeo Parcellation map on a sample human 32k fsLR surface and **B)** associated functional network node distribution across the gray matter. Within the human parcellation, the functional network distributions for the seven networks are relatively evenly distributed, ranging from 7-19% of the gray matter surface. The Default Network is associated with the highest number of nodes within the 32k fsLR human surface, accounting for 19% of assigned nodes. The Limbic Network is the lowest assigned network, accounting for 7% of the assigned nodes within the gray matter surface. **C)** Yeo Parcellation map aligned to a sample macaque 32k fsLR surface and **D)** associated functional network node distribution. Dominant networks are the Visual Network (26%) and the Somatomotor Network (16%). The functional networks primarily thought to be associated with higher level cognitive processing, the Default Network and Frontoparietal Network, are more defined in the Human parcellation compared to the NHP parcellation, 19% vs. 12% and 12% vs. 7% surface area, respectively.

### Individual Specific TMS Functional Network Analysis

To assign the functional connectivity maps to the Yeo-7 networks, we first applied a sparsity threshold to keep only the top 1% correlation values for each functional connectivity map. Other sparsity thresholds lead to similar network activation patterns (**Supplementary Figure 2, Supplementary Figure 3**). We binarized the functional connectivity maps and calculated the percentage of overlap between the binarized maps for each stimulation configuration to each Yeo parcellation network. Other assignment metrics lead to similar network activation patterns (**Supplementary Figure 4**). The network activation evaluation for all participants in the human and macaque datasets was conducted in separate analyses.

### Intra-Species TMS Functional Network Analysis

To reduce the complexity of the group analysis, we assigned the TMS locations across participants to nine zones (Z1-Z9) oriented in a 3×3 grid. A single stimulation zone accounted for four coil locations and 12 coil orientations per participant (**Figure 3A**). Using these nine stimulation zones, we analyzed the overlap of the TMS activation profile with the Yeo-7 human and macaque parcellations. We identified the zone’s predicted functional network based on the highest overlap with the Yeo-7 networks. In a secondary analysis, we analyzed the effect of TMS coil orientation within each stimulation zone separately. We reduced the simulated coil orientations into groups of 45-degree steps to account for variations in TMS targeting, resulting in four coil orientations per coil location. The highest overlap coefficients across each zone’s orientation windows were calculated to identify the effect of coil orientation on the predicted functional networks.

**Figure 3.**
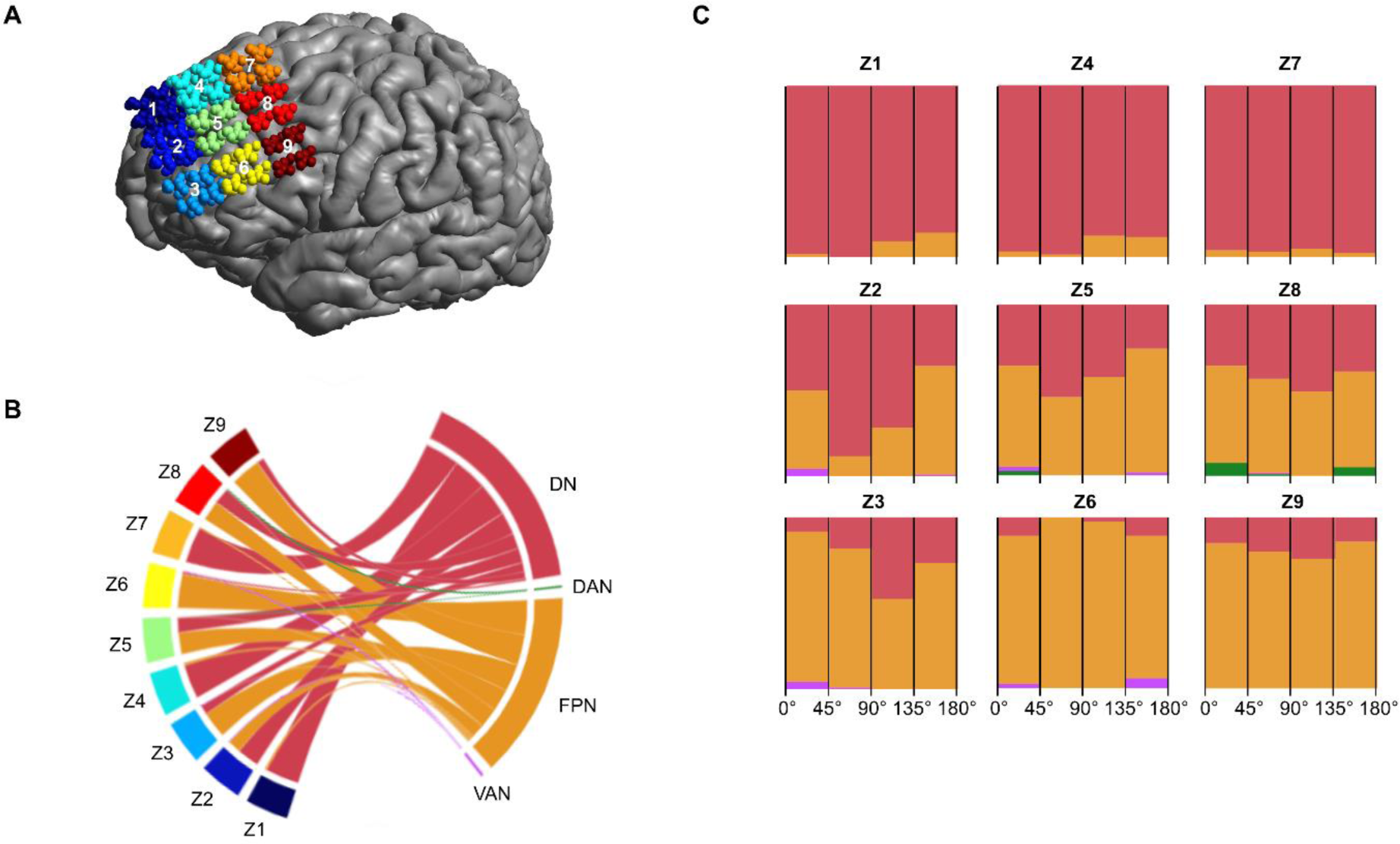
Overlap of TMS functional networks with Yeo Networks in Humans. **A)** We investigated the effect of coil location across all participants on a 3×3 zoned grid (Z1-Z9), with each grid zone representing four simulation locations per participant. **B)** The relative overlap of the stimulation zone with Yeo functional parcellations is determined by summing the functional connectivity maps generated for each TMS coil configuration (coil location and orientation). We identify the highest overlapping functional networks across each stimulation zone. For humans, the Default Network exhibits the highest activation within the medial zones (Z1, Z4, Z5), while the Frontoparietal Network is highly activated within the lateral zones (Z3, Z6, Z9). A transition zone (Z2, Z5, Z8) has contributions from both networks. **C.** To examine the effect of coil orientation, we condensed the 12 coil orientations into four groups, each representing a 45-degree range, within each stimulation zone. TMS network activation is largely independent of orientation and is primarily influenced by the location-based activation of the Default and Frontoparietal Networks. In the transition zones (Z2, Z5, Z8), network activation differs between Default and Frontoparietal Networks depending on coil orientation.

### Inter-Species TMS Functional Network Analysis

In order to explain why specific functional networks were more activated than others we identified areas that were preferentially stimulated by TMS across all grid locations in the prefrontal cortex. We calculated an activation index which highlights brain regions most often targeted by TMS across all simulations. For this we identified the percentage of nodes within GM that were above the 50% maximum electric field threshold across all 432 simulations. To assess the impact of brain gyrification on found results, we repeated the same analysis for a homogeneous conductivity model (Supplementary Figure 5). The activation index was calculated for both species separately.

## Results

### Overlap of TMS Activation with Established Functional Networks

We first investigated the TMS functional networks for the human participants. We found that the Frontoparietal Network (FPN) and Default Network (DN) dominated the predicted network activation across stimulation locations (**Figure 3A, B**). Specifically, the FPN was most prominent in the lateral zones (Z3, Z6, Z9), while the DN was more activated in the medial zones (Z1, Z4, Z7). There was a transition between the activation of these two functional networks in the middle zones (Z2, Z5, Z8). To analyze the effect of coil orientation on network activation, we grouped the coil orientations into four groups, representing coil orientations ranges of 45 degrees (**Figure 3C**). We found that in the transition zone functional networks differed across coil orientations, with the FPN and DN roughly equally activated. These findings align with previously modeled TMS functional networks in prefrontal brain regions (Opitz et al., 2016).

Next, we investigated TMS functional networks in macaques. Here, the FPN dominates the predicted network activation within the frontal zones (Z1, Z2, Z3, Z4, Z5) (**Figure 4A, B**). The posterior zones (Z6, Z7, Z8, Z9) are a transition zone between the FPN and the Somatomotor Network (SN). We grouped the coil orientations while analyzing the effect of coil orientation on network activation in the same manner as for the human participants. Here, we found the presence of secondary networks that were activated only few times (DN, Dorsal Attention Network (DAN), and Ventral Attention Network (VAN)) throughout all zones, irrespective of coil orientation (**Figure 4C**).

**Figure 4.**
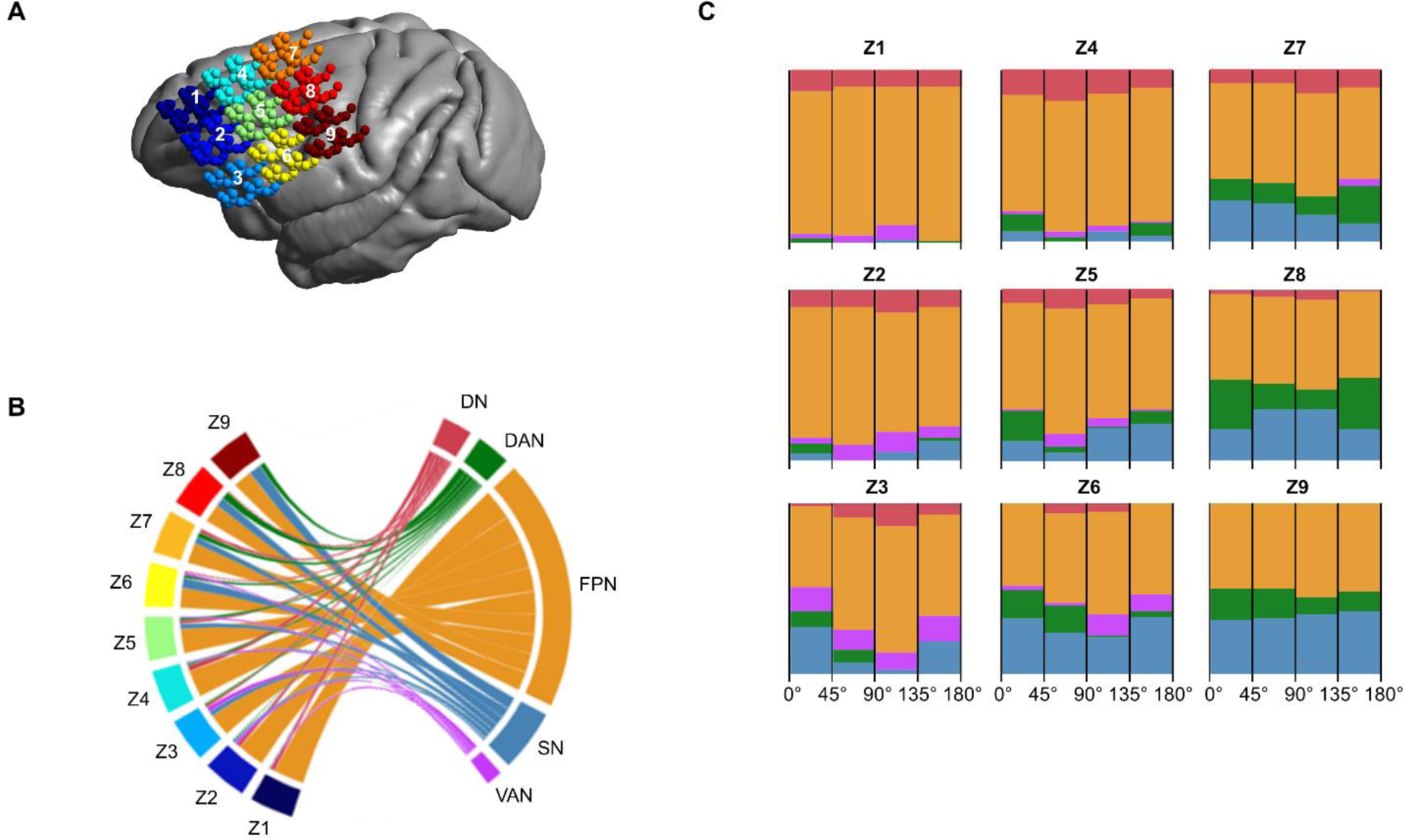
Overlap of TMS functional networks with Yeo Networks in Macaques. **A)** We investigated the effect of coil location across all participants on a 3×3 zoned grid, with each grid section representing four simulation locations per individual. **B)** The relative overlap of the stimulation zone with Yeo functional parcellations is determined by summing the functional connectivity maps generated for each stimulation coil configuration (coil location and orientation). The highest overlapping functional networks are aggregated within each zone. The Frontoparietal Network activation is most prominent within the frontal zones (Z1, Z2, Z3, Z4, Z5), with some default network activation in the most medial locations. The posterior zones (Z6, Z7, Z8, Z9) act as a transition zone between Frontoparietal Network activation and other networks (Somatomotor, Dorsal Attention, Ventral Attention). **C)** To analyze the effect of coil orientation, the 12 coil orientations were reduced to four groups (encompassing 45 degrees) within each stimulation zone. Within the NHP dataset, the TMS network activation is largely orientation independent.

### Cross-Species Comparison of Functional Network Activation

We investigated which brain regions exhibited the highest electric field activation across all 432 simulations to investigate the difference in functional network activation across species. The activation index represented the frequency of electric field activation within a specific brain region (> 50% of Emax) across all simulations. For the human models, we found a uniform spread across the prefrontal cortex with maximum activation regions of 91% across all simulations within the transitional zone (**Figure 5A**). The two primary functional networks that are activated by TMS in the human prefrontal cortex are the DN and the FPN, being targeted in 53% and 46% of all stimulation conditions, respectively (**Figure 5B**). In contrast, other functional networks, including DAN and VAN, exhibit less frequent activation.

**Figure 5.**
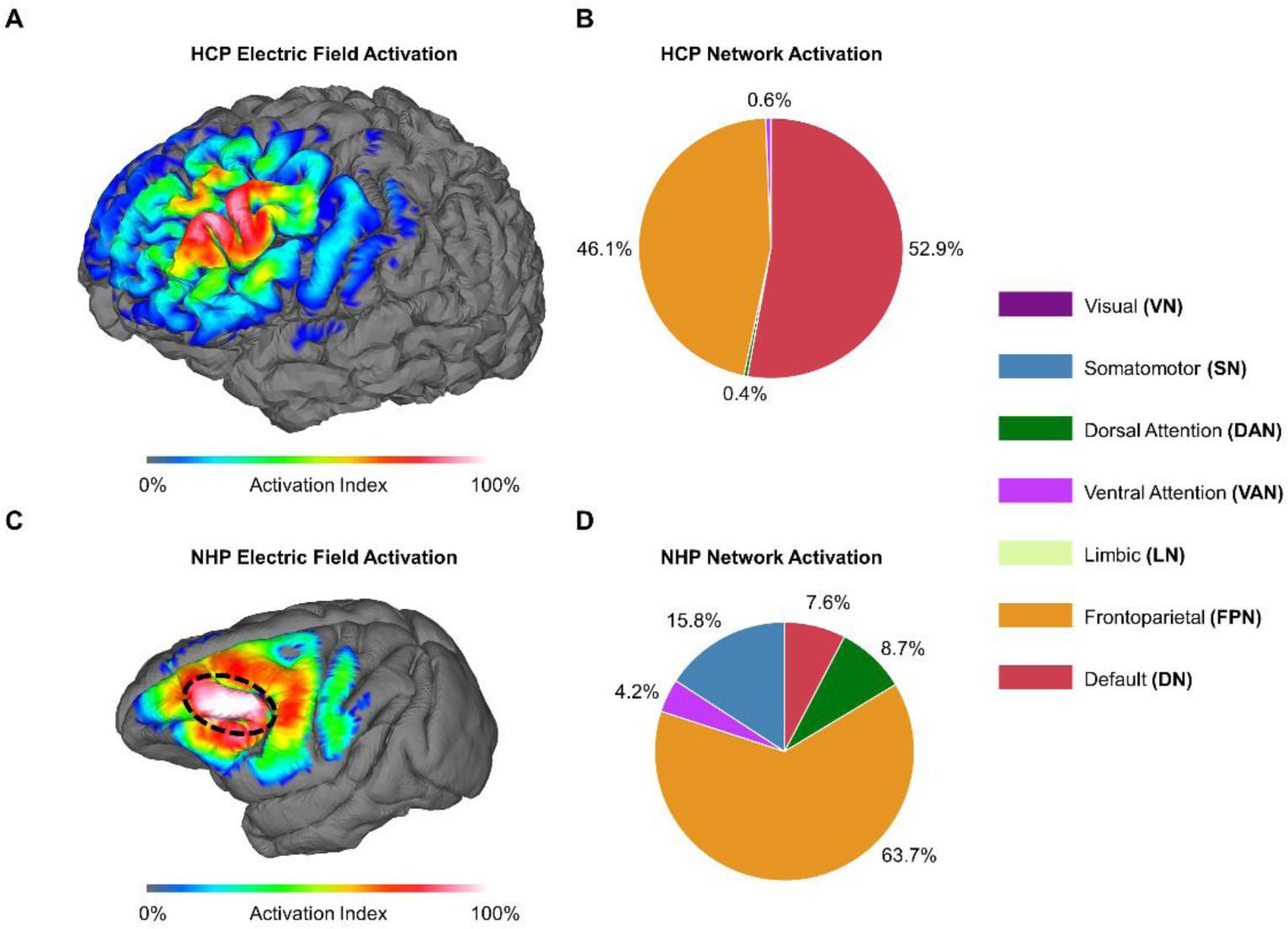
Comparative functional network activation across humans and macaques. **A)** The electric field activation index is relatively distributed across the stimulation grid across all simulation configurations within the human models. The activation index is the highest in the center of the grid (91%), aligning with the evaluated stimulation transition zone. **B)** In the human models, the DN and FPN are activated by 53% and 46% of the stimulation conditions, respectively. Other functional networks, such as DAN and VAN, are activated, but to a lesser extent. **C)** In the NHP models, the electric field activation pattern is concentrated along the distinct gyral fold in the prefrontal cortex. The activation index, summed across all simulations, demonstrates a preferential activation of this area and less electric field dispersion when compared to the observed pattern in the human models. In the Yeo NHP parcellation, this gyral fold is associated primarily with the Frontoparietal Network. **D)** The Frontoparietal Network is the most prominent functional network targeted by prefrontal TMS, activated by 64% of all stimulation conditions. The Somatomotor Network (SN) is activated by 16% of the stimulation conditions. To a lesser extent, other functional networks, DAN, VAN, and DN are activated.

In all simulation conditions within the NHP model, the electric field strength concentrates along the distinct gyral fold (i.e., area 8a and 46d) in the prefrontal cortex (**Figure 5C**). This brain region is activated in almost all conditions (close to 100%). This is likely due to the surrounding sulci (i.e. principle sulcus and arcuate sulcus) leading to increased electric field strengths at the CSF-GM interface (Thielscher et al., 2011). In a homogenous conductivity model, the electric field strength is more evenly distributed (**Supplementary Figure 5**). Within the Yeo primate parcellation, this gyral fold is primarily associated with the FPN. Thus, the FPN emerges as the dominant target, being activated in 64% of all stimulation conditions (**Figure 5D**). In addition, we found comparatively higher activation of other functional networks in the NHP model compared to the human model. The NHP models show activation of the SN (16%), DAN (9%), VAN (4%) and DN (8%).

## Discussion

In this study we compared functional networks in the prefrontal cortex targeted by TMS across humans and macaques. We investigated different TMS coil locations and orientations and identified the affected functional networks in humans and macaques. In line with previous work, we found two dominant networks are activated in humans: The frontoparietal network (FPN) and the default network (DN) (Opitz et al., 2016). Specifically, we found 1) a dominant DN activation in a medial, orientation-insensitive zone, 2) a dominant FPN activation in a lateral orientation-insensitive zone, and 3) a transition zone that allowed for activation of either the DN or the FPN depending on TMS coil orientation. In the NHP analysis, we found that the FPN was predominantly activated by the anterior stimulation zones, which was coil orientation-insensitive. In more posterior stimulation zones, we found that the FPN remained the dominant targeted network, but depending on coil orientation, targeting of the somatomotor network or dorsal attention network co-occurred.

An important distinction between human and NHP TMS functional networks is the ability to target the DN. We found that in macaques, targeting the corresponding DN in the prefrontal lobe is notably more challenging than in humans. One reason for this observation is the difference in brain gyrification across species. Compared to humans, the macaque NHP cortical surface is less gyrated (Hofman, 2014; Van Essen and Dierker, 2007). In the prefrontal cortex, the human brain is highly gyrated, while the macaque brain has one prominent gyrus. The gyrification pattern is important because it affects the TMS-induced electric field. TMS electric fields are enhanced in GM when currents are crossing the CSF-GM interface (Miranda et al., 2007; Thielscher et al., 2011). The anatomical features, including gyri, of the brain affect which regions get preferentially stimulated across the investigated TMS coil locations (**Figure 5**). In humans, we found strong activation within the DN-FPN transition zone and evenly distributed activation in each network-specific (DN, FPN) stimulation zone. In comparison, we found preferential stimulation of the dominant cortical fold primarily corresponding to the FPN in macaques. Thus, differences in TMS functional network activation across species are affected by differences in brain gyrification.

The NHP and human network activation profiles exhibit differences in targeting secondary functional networks. In the human models, the investigated TMS coil locations equally target two distinct networks (DN and FPN). In contrast, in the NHP models, there is a preferential activation of the FPN network, but we see increased secondary network targeting. These secondary networks include the dorsal attention network, somatomotor network, ventral attention network, and DN. The increased activity in secondary networks can be attributed to the extent of electric field distribution on the NHP cortical surface. Notably, the cortical gray matter of the human prefrontal cortex exceeds that of the macaques by up to 1.9-fold (Donahue et al., 2018). Additionally, the smaller brain size in NHPs results in a larger electric field distribution relative to surface area (Alekseichuk et al., 2019), leading to the incidental activation of other networks outside the targeted ones. The prefrontal cortex in humans is remarkably expanded, resulting in pronounced connections to multimodal areas and greater network modularity (Garin et al., 2022; Liu et al., 2019; Mantini et al., 2011). DN stands out as the network exhibiting the most significant differences in both structure and function between humans and NHPs (Xu et al., 2020). Although DN appears to be present in NHPs in the medial frontal and the posterior cingulate cortex, the lateral prefrontal cortex in macaque likely plays a role in attention and executive functions (Bahmani et al., 2019; Bullock et al., 2017; Petrides et al., 2012). These differences underscore the translational challenges between human and macaque network targeting.

The comparison of TMS functional networks between humans and NHPs did not reveal a direct one-to-one mapping across species in the prefrontal cortex. The distinct differences in each species’ anatomical and functional brain organization lead to different functional networks targeted by TMS, highlighting the challenges of cross-species translation. Our comprehensive study of TMS functional network targeting can thus inform future translational efforts. Prefrontal TMS is relevant in the treatment of various neuropsychiatric disorders such as depression. Brain stimulation therapies targeting functional networks within this region are a promising tool to modulate neural circuits affected by specific disorders (Siddiqi et al., 2020). NHPs offer a unique opportunity to validate and refine TMS network targeting strategies. Our cross-species network comparison can guide these efforts to help improve translational efforts in NHPs. Future experimental work is needed to validate the predictions made by our network targeting approach in both humans and NHPs.

## Acknowledgements

We thank Kathleen Mantell and Jonna Rotteveel for their help with the manual segmentation of the non-human primate models. This work was supported by RF1MH117428, RF1MH128696, R01NS109498, and a National Science Foundation Graduate Research Fellowship.

## Authors Contributions

T.B.: Conceptualization, Methodology, Software, Formal Analysis, Data Curation, Writing – Original Draft, Funding Acquisition; A.O.: Conceptualization, Methodology, Supervision, Writing – Review & Editing, Funding Acquisition; T.X.: Conceptualization, Methodology, Data Curation, Writing – Review & Editing, Funding Acquisition.

## References

Alekseichuk, I., Mantell, K., Shirinpour, S., Opitz, A., 2019. Comparative modeling of transcranial magnetic and electric stimulation in mouse, monkey, and human. NeuroImage 194, 136–148. 10.1016/j.neuroimage.2019.03.044

Amiez, C., Kostopoulos, P., Champod, A.-S., Petrides, M., 2006. Local Morphology Predicts Functional Organization of the Dorsal Premotor Region in the Human Brain. J. Neurosci. 26, 2724–2731. 10.1523/JNEUROSCI.4739-05.2006

Ardesch, D.J., Scholtens, L.H., Li, L., Preuss, T.M., Rilling, J.K., van den Heuvel, M.P., 2019. Evolutionary expansion of connectivity between multimodal association areas in the human brain compared with chimpanzees. Proc. Natl. Acad. Sci. 116, 7101–7106. 10.1073/pnas.1818512116

Avery, D.H., Holtzheimer, P.E., Fawaz, W., Russo, J., Neumaier, J., Dunner, D.L., Haynor, D.R., Claypoole, K.H., Wajdik, C., Roy-Byrne, P., 2006. A Controlled Study of Repetitive Transcranial Magnetic Stimulation in Medication-Resistant Major Depression. Biol. Psychiatry 59, 187–194. 10.1016/j.biopsych.2005.07.003

Bahmani, Z., Clark, K., Merrikhi, Y., Mueller, A., Pettine, W., Isabel Vanegas, M., Moore, T., Noudoost, B., 2019. Prefrontal Contributions to Attention and Working Memory, in: Hodgson, T. (Ed.), Processes of Visuospatial Attention and Working Memory, Current Topics in Behavioral Neurosciences. Springer International Publishing, Cham, pp. 129–153. 10.1007/7854_2018_74

Bullock, K.R., Pieper, F., Sachs, A.J., Martinez-Trujillo, J.C., 2017. Visual and presaccadic activity in area 8Ar of the macaque monkey lateral prefrontal cortex. J. Neurophysiol. 118, 15–28. 10.1152/jn.00278.2016

de Lima-Pardini, A.C., Mikhail, Y., Dominguez-Vargas, A.-U., Dancause, N., Scott, S.H., 2023. Transcranial magnetic stimulation in non-human primates: A systematic review. Neurosci. Biobehav. Rev. 152, 105273. 10.1016/j.neubiorev.2023.105273

Deng, Z.-D., Lisanby, S.H., Peterchev, A.V., 2013. Electric field depth–focality tradeoff in transcranial magnetic stimulation: Simulation comparison of 50 coil designs. Brain Stimulat. 6, 1–13. 10.1016/j.brs.2012.02.005

Donahue, C.J., Glasser, M.F., Preuss, T.M., Rilling, J.K., Van Essen, D.C., 2018. Quantitative assessment of prefrontal cortex in humans relative to nonhuman primates. Proc. Natl. Acad. Sci. 115, E5183– E5192. 10.1073/pnas.1721653115

Fox, M.D., Halko, M.A., Eldaief, M.C., Pascual-Leone, A., 2012. Measuring and manipulating brain connectivity with resting state functional connectivity magnetic resonance imaging (fcMRI) and transcranial magnetic stimulation (TMS). Neuroimage 62, 2232–2243. 10.1016/j.neuroimage.2012.03.035

Garin, C.M., Hori, Y., Everling, S., Whitlow, C.T., Calabro, F.J., Luna, B., Froesel, M., Gacoin, M., Ben Hamed, S., Dhenain, M., Constantinidis, C., 2022. An evolutionary gap in primate default mode network organization. Cell Rep. 39, 110669. 10.1016/j.celrep.2022.110669

George, M.S., Lisanby, S.H., Avery, D., McDonald, W.M., Durkalski, V., Pavlicova, M., Anderson, B., Nahas, Z., Bulow, P., Zarkowski, P., Holtzheimer, P.E., III, Schwartz, T., Sackeim, H.A., 2010. Daily Left Prefrontal Transcranial Magnetic Stimulation Therapy for Major Depressive Disorder: A Sham-Controlled Randomized Trial. Arch. Gen. Psychiatry 67, 507–516. 10.1001/archgenpsychiatry.2010.46

George, M.S., Wassermann, E.M., Williams, W.A., Callahan, A., Ketter, T.A., Basser, P., Hallett, M., Post, R.M., 1995. Daily repetitive transcranial magnetic stimulation (rTMS) improves mood in depression. Neuroreport 6, 1853–1856. 10.1097/00001756-199510020-00008

Glasser, M.F., Sotiropoulos, S.N., Wilson, J.A., Coalson, T.S., Fischl, B., Andersson, J.L., Xu, J., Jbabdi, S., Webster, M., Polimeni, J.R., Van Essen, D.C., Jenkinson, M., WU-Minn HCP Consortium, 2013. The minimal preprocessing pipelines for the Human Connectome Project. NeuroImage 80, 105–124. 10.1016/j.neuroimage.2013.04.127

Goulas, A., Uylings, H.B.M., Stiers, P., 2012. Unravelling the Intrinsic Functional Organization of the Human Lateral Frontal Cortex: A Parcellation Scheme Based on Resting State fMRI. J. Neurosci. 32, 10238–10252. 10.1523/JNEUROSCI.5852-11.2012

Gratton, C., Kraus, B.T., Greene, D.J., Gordon, E.M., Laumann, T.O., Nelson, S.M., Dosenbach, N.U.F., Petersen, S.E., 2020. Defining Individual-Specific Functional Neuroanatomy for Precision Psychiatry. Biol. Psychiatry 88, 28–39. 10.1016/j.biopsych.2019.10.026

Hamada, M., Murase, N., Hasan, A., Balaratnam, M., Rothwell, J.C., 2013. The role of interneuron networks in driving human motor cortical plasticity. Cereb. Cortex N. Y. N 1991 23, 1593–1605. 10.1093/cercor/bhs147

Hanlon, C.A., Czoty, P.W., Smith, H.R., Epperly, P.M., Galbo, L.K., 2021. Cortical excitability in a nonhuman primate model of TMS. Brain Stimulat. 14, 19–21. 10.1016/j.brs.2020.10.008

Herbsman, T., Avery, D., Ramsey, D., Holtzheimer, P., Wadjik, C., Hardaway, F., Haynor, D., George, M.S., Nahas, Z., 2009. More Lateral and Anterior Prefrontal Coil Location Is Associated with Better Repetitive Transcranial Magnetic Stimulation Antidepressant Response. Biol. Psychiatry, Medical Consequences and Contributions to Depression 66, 509–515. 10.1016/j.biopsych.2009.04.034

Herwig, U., Padberg, F., Unger, J., Spitzer, M., Schönfeldt-Lecuona, C., 2001. Transcranial magnetic stimulation in therapy studies: examination of the reliability of “standard” coil positioning by neuronavigation. Biol. Psychiatry 50, 58–61. 10.1016/S0006-3223(01)01153-2

Hofman, M., 2014. Evolution of the human brain: when bigger is better. Front. Neuroanat. 8.

Laumann, T.O., Gordon, E.M., Adeyemo, B., Snyder, A.Z., Joo, S.J., Chen, M.-Y., Gilmore, A.W., McDermott, K.B., Nelson, S.M., Dosenbach, N.U.F., Schlaggar, B.L., Mumford, J.A., Poldrack, R.A., Petersen, S.E., 2015. Functional system and areal organization of a highly sampled individual human brain. Neuron 87, 657–670. 10.1016/j.neuron.2015.06.037

Lear, A., Baker, S.N., Clarke, H.F., Roberts, A.C., Schmid, M.C., Jarrett, W., 2022. Understanding them to understand ourselves: The importance of NHP research for translational neuroscience. Curr. Res. Neurobiol. 3, 100049. 10.1016/j.crneur.2022.100049

Lefaucheur, J.-P., André-Obadia, N., Antal, A., Ayache, S.S., Baeken, C., Benninger, D.H., Cantello, R.M., Cincotta, M., de Carvalho, M., De Ridder, D., Devanne, H., Di Lazzaro, V., Filipović, S.R., Hummel, F.C., Jääskeläinen, S.K., Kimiskidis, V.K., Koch, G., Langguth, B., Nyffeler, T., Oliviero, A., Padberg, F., Poulet, E., Rossi, S., Rossini, P.M., Rothwell, J.C., Schönfeldt-Lecuona, C., Siebner, H.R., Slotema, C.W., Stagg, C.J., Valls-Sole, J., Ziemann, U., Paulus, W., Garcia-Larrea, L., 2014. Evidence-based guidelines on the therapeutic use of repetitive transcranial magnetic stimulation (rTMS). Clin. Neurophysiol. 125, 2150–2206. 10.1016/j.clinph.2014.05.021

Liu, C., Yen, C.C.-C., Szczupak, D., Ye, F.Q., Leopold, D.A., Silva, A.C., 2019. Anatomical and functional investigation of the marmoset default mode network. Nat. Commun. 10, 1975. 10.1038/s41467-019-09813-7

López-Alonso, V., Cheeran, B., Río-Rodríguez, D., Fernández-del-Olmo, M., 2014. Inter-individual Variability in Response to Non-invasive Brain Stimulation Paradigms. Brain Stimulat. 10.1016/j.brs.2014.02.004

Mantini, D., Gerits, A., Nelissen, K., Durand, J.-B., Joly, O., Simone, L., Sawamura, H., Wardak, C., Orban, G.A., Buckner, R.L., Vanduffel, W., 2011. Default Mode of Brain Function in Monkeys. J. Neurosci. 31, 12954–12962. 10.1523/JNEUROSCI.2318-11.2011

Margulies, D.S., Petrides, M., 2013. Distinct Parietal and Temporal Connectivity Profiles of Ventrolateral Frontal Areas Involved in Language Production. J. Neurosci. 33, 16846–16852. 10.1523/JNEUROSCI.2259-13.2013

Mars, R.B., Sotiropoulos, S.N., Passingham, R.E., Sallet, J., Verhagen, L., Khrapitchev, A.A., Sibson, N., Jbabdi, S., 2018. Whole brain comparative anatomy using connectivity blueprints. eLife 7, e35237. 10.7554/eLife.35237

Milham, M.P., Ai, L., Koo, B., Xu, T., Amiez, C., Balezeau, F., Baxter, M.G., Blezer, E.L.A., Brochier, T., Chen, A., Croxson, P.L., Damatac, C.G., Dehaene, S., Everling, S., Fair, D.A., Fleysher, L., Freiwald, W., Froudist-Walsh, S., Griffiths, T.D., Guedj, C., Hadj-Bouziane, F., Ben Hamed, S., Harel, N., Hiba, B., Jarraya, B., Jung, B., Kastner, S., Klink, P.C., Kwok, S.C., Laland, K.N., Leopold, D.A., Lindenfors, P., Mars, R.B., Menon, R.S., Messinger, A., Meunier, M., Mok, K., Morrison, J.H., Nacef, J., Nagy, J., Rios, M.O., Petkov, C.I., Pinsk, M., Poirier, C., Procyk, E., Rajimehr, R., Reader, S.M., Roelfsema, P.R., Rudko, D.A., Rushworth, M.F.S., Russ, B.E., Sallet, J., Schmid, M.C., Schwiedrzik, C.M., Seidlitz, J., Sein, J., Shmuel, A., Sullivan, E.L., Ungerleider, L., Thiele, A., Todorov, O.S., Tsao, D., Wang, Z., Wilson, C.R.E., Yacoub, E., Ye, F.Q., Zarco, W., Zhou, Y., Margulies, D.S., Schroeder, C.E., 2018. An Open Resource for Non-human Primate Imaging. Neuron 100, 61–74.e2. 10.1016/j.neuron.2018.08.039

Miranda, P.C., Correia, L., Salvador, R., Basser, P.J., 2007. Tissue heterogeneity as a mechanism for localized neural stimulation by applied electric fields. Phys. Med. Biol. 52, 5603–17. 10.1088/0031-9155/52/18/009

Mueller, J.K., Grigsby, E.M., Prevosto, V., Petraglia, F.W., Rao, H., Deng, Z.-D., Peterchev, A.V., Sommer, M.A., Egner, T., Platt, M.L., Grill, W.M., 2014. Simultaneous transcranial magnetic stimulation and single-neuron recording in alert non-human primates. Nat. Neurosci. 17, 1130– 1136. 10.1038/nn.3751

Noonan, M.P., Sallet, J., Mars, R.B., Neubert, F.X., O’Reilly, J.X., Andersson, J.L., Mitchell, A.S., Bell, A.H., Miller, K.L., Rushworth, M.F.S., 2014. A Neural Circuit Covarying with Social Hierarchy in Macaques. PLOS Biol. 12, e1001940. 10.1371/journal.pbio.1001940

Oathes, D.J., Zimmerman, J.P., Duprat, R., Japp, S.S., Scully, M., Rosenberg, B.M., Flounders, M.W., Long, H., Deluisi, J.A., Elliott, M., Shandler, G., Shinohara, R.T., Linn, K.A., 2021. Resting fMRI guided TMS results in subcortical and brain network modulation indexed by interleaved TMS/fMRI. Exp. Brain Res. 239, 1165–1178. 10.1007/s00221-021-06036-5

Opitz, A., Fox, M.D., Craddock, R.C., Colcombe, S., Milham, M.P., 2016. An integrated framework for targeting functional networks via transcranial magnetic stimulation. NeuroImage 127, 86–96. 10.1016/j.neuroimage.2015.11.040

Opitz, A., Windhoff, M., Heidemann, R.M., Turner, R., Thielscher, A., 2011. How the brain tissue shapes the electric field induced by transcranial magnetic stimulation. NeuroImage 58. 10.1016/j.neuroimage.2011.06.069

Pascual-Leone, A., Rubio, B., Pallardó, F., Catalá, M.D., 1996. Rapid-rate transcranial magnetic stimulation of left dorsolateral prefrontal cortex in drug-resistant depression. The Lancet 348, 233–237. 10.1016/S0140-6736(96)01219-6

Perera, N.D., Alekseichuk, I., Shirinpour, S., Wischnewski, M., Linn, G., Masiello, K., Butler, B., Russ, B.E., Schroeder, C.E., Falchier, A., Opitz, A., 2023. Dissociation of Centrally and Peripherally Induced Transcranial Magnetic Stimulation Effects in Nonhuman Primates. J. Neurosci. 10.1523/JNEUROSCI.1016-23.2023

Petrides, M., Tomaiuolo, F., Yeterian, E.H., Pandya, D.N., 2012. The prefrontal cortex: Comparative architectonic organization in the human and the macaque monkey brains. Cortex, Frontal lobes 48, 46–57. 10.1016/j.cortex.2011.07.002

Power, J.D., Cohen, A.L., Nelson, S.M., Wig, G.S., Barnes, K.A., Church, J.A., Vogel, A.C., Laumann, T.O., Miezin, F.M., Schlaggar, B.L., Petersen, S.E., 2011. Functional network organization of the human brain. Neuron 72, 665–678. 10.1016/j.neuron.2011.09.006

Rizvi, S., Khan, A.M., 2019. Use of Transcranial Magnetic Stimulation for Depression. Cureus 11. 10.7759/cureus.4736

Romero, M.C., Davare, M., Armendariz, M., Janssen, P., 2019. Neural basis of Transcranial Magnetic Stimulation at the single-cell Level. Nat. Commun. 405753–405753. 10.1101/405753

Rossi, S., Hallett, M., Rossini, P.M., Pascual-Leone, A., Safety of TMS Consensus Group, 2009. Safety, ethical considerations, and application guidelines for the use of transcranial magnetic stimulation in clinical practice and research. Clin. Neurophysiol. Off. J. Int. Fed. Clin. Neurophysiol. 120, 2008–2039. 10.1016/j.clinph.2009.08.016

Rusinkiewicz, S., Levoy, M., 2001. Efficient variants of the ICP algorithm, in: Proceedings Third International Conference on 3-D Digital Imaging and Modeling. Presented at the Proceedings Third International Conference on 3-D Digital Imaging and Modeling, pp. 145–152. 10.1109/IM.2001.924423

Schutter, D.J.L.G., 2009. Antidepressant efficacy of high-frequency transcranial magnetic stimulation over the left dorsolateral prefrontal cortex in double-blind sham-controlled designs: a meta-analysis. Psychol. Med. 39, 65–75. 10.1017/S0033291708003462

Siddiqi, S.H., Taylor, S.F., Cooke, D., Pascual-Leone, A., George, M.S., Fox, M.D., 2020. Distinct symptom-specific treatment targets for circuit-based neuromodulation. Am. J. Psychiatry 177, 435–446. 10.1176/appi.ajp.2019.19090915

Smith, Stephen M, Andersson, J., Auerbach, E.J., Beckmann, C.F., Bijsterbosch, J., Douaud, G., Duff, E., Feinberg, D.A., Griffanti, L., Harms, M.P., Kelly, M., Laumann, T., Miller, K.L., Moeller, S., Petersen, S., Power, J., Salimi-Khorshidi, G., Snyder, A.Z., Vu, A., Woolrich, M.W., Xu, J., Yacoub, E., Ugurbil, K., Van Essen, D., Glasser, M.F., 2013. Resting-state fMRI in the Human Connectome Project. NeuroImage 80, 144–168. 10.1016/j.neuroimage.2013.05.039

Smith, Stephen M., Beckmann, C.F., Andersson, J., Auerbach, E.J., Bijsterbosch, J., Douaud, G., Duff, E., Feinberg, D. a., Griffanti, L., Harms, M.P., Kelly, M., Laumann, T., Miller, K.L., Moeller, S., Petersen, S., Power, J., Salimi-Khorshidi, G., Snyder, A.Z., Vu, A.T., Woolrich, M.W., Xu, J., Yacoub, E., Uğurbil, K., Van Essen, D.C., Glasser, M.F., Uǧurbil, K., Van Essen, D.C., Glasser, M.F., Uğurbil, K., Van Essen, D.C., Glasser, M.F., 2013. Resting-state fMRI in the Human Connectome Project. NeuroImage 80, 144–168. 10.1016/j.neuroimage.2013.05.039

Thielscher, A., Antunes, A., Saturnino, G.B., 2015. Field modeling for transcranial magnetic stimulation: A useful tool to understand the physiological effects of TMS? Annu. Int. Conf. IEEE Eng. Med. Biol. Soc. IEEE Eng. Med. Biol. Soc. Annu. Int. Conf. 2015, 222–225. 10.1109/EMBC.2015.7318340

Thielscher, A., Opitz, A., Windhoff, M., 2011. Impact of the gyral geometry on the electric field induced by transcranial magnetic stimulation. NeuroImage 54, 234–243. 10.1016/j.neuroimage.2010.07.061

van den Heuvel, M.P., Ardesch, D.J., Scholtens, L.H., de Lange, S.C., van Haren, N.E.M., Sommer, I.E.C., Dannlowski, U., Repple, J., Preuss, T.M., Hopkins, W.D., Rilling, J.K., 2023. Human and chimpanzee shared and divergent neurobiological systems for general and specific cognitive brain functions. Proc. Natl. Acad. Sci. 120, e2218565120. 10.1073/pnas.2218565120

Van Essen, D.C., 2004. Surface-based approaches to spatial localization and registration in primate cerebral cortex. NeuroImage 23 Suppl 1, S97–107. 10.1016/j.neuroimage.2004.07.024

Van Essen, D.C., Dierker, D.L., 2007. Surface-Based and Probabilistic Atlases of Primate Cerebral Cortex. Neuron 56, 209–225. 10.1016/j.neuron.2007.10.015

Van Essen, D.C., Ugurbil, K., Auerbach, E., Barch, D., Behrens, T.E.J., Bucholz, R., Chang, A., Chen, L., Corbetta, M., Curtiss, S.W., Della Penna, S., Feinberg, D., Glasser, M.F., Harel, N., Heath, A.C., Larson-Prior, L., Marcus, D., Michalareas, G., Moeller, S., Oostenveld, R., Petersen, S.E., Prior, F., Schlaggar, B.L., Smith, S.M., Snyder, A.Z., Xu, J., Yacoub, E., WU-Minn HCP Consortium, 2012. The Human Connectome Project: a data acquisition perspective. NeuroImage 62, 2222– 2231. 10.1016/j.neuroimage.2012.02.018

Windhoff, M., Opitz, A., Thielscher, A., 2013. Electric field calculations in brain stimulation based on finite elements: An optimized processing pipeline for the generation and usage of accurate individual head models. Hum. Brain Mapp. 34, 923–35. 10.1002/hbm.21479

WU_Minn, H.C.P., 2017. 1200 Subject data release reference manual [WWW Document]. URL https://www.humanconnectome.org/storage/app/media/documentation/s1200/HCP_S1200_Release_Reference_Manual.pdf (accessed 6.30.22).

Xu, T., Falchier, A., Sullivan, E.L., Linn, G., Ramirez, J.S.B., Ross, D., Feczko, E., Opitz, A., Bagley, J., Sturgeon, D., Earl, E., Miranda-Domínguez, O., Perrone, A., Craddock, R.C., Schroeder, C.E., Colcombe, S., Fair, D.A., Milham, M.P., 2018. Delineating the Macroscale Areal Organization of the Macaque Cortex In Vivo. Cell Rep. 23, 429–441. 10.1016/j.celrep.2018.03.049

Xu, T., Nenning, K.-H., Schwartz, E., Hong, S.-J., Vogelstein, J.T., Goulas, A., Fair, D.A., Schroeder, C.E., Margulies, D.S., Smallwood, J., Milham, M.P., Langs, G., 2020. Cross-species functional alignment reveals evolutionary hierarchy within the connectome. NeuroImage 223, 117346. 10.1016/j.neuroimage.2020.117346

Xu, T., Sturgeon, D., Ramirez, J.S.B., Froudist-Walsh, S., Margulies, D.S., Schroeder, C.E., Fair, D.A., Milham, M.P., 2019. Interindividual Variability of Functional Connectivity in Awake and Anesthetized Rhesus Macaque Monkeys. Biol. Psychiatry Cogn. Neurosci. Neuroimaging 4, 543–553. 10.1016/j.bpsc.2019.02.005

Xu, T., Yang, Z., Jiang, L., Xing, X.-X., Zuo, X.-N., 2015. A Connectome Computation System for discovery science of brain. Sci. Bull. 60, 86–95. 10.1007/s11434-014-0698-3

Yeo, B.T.T., Krienen, F.M., Sepulcre, J., Sabuncu, M.R., Lashkari, D., Hollinshead, M., Roffman, J.L., Smoller, J.W., Zöllei, L., Polimeni, J.R., Fischl, B., Liu, H., Buckner, R.L., 2011. The organization of the human cerebral cortex estimated by intrinsic functional connectivity. J. Neurophysiol. 106, 1125–1165. 10.1152/jn.00338.2011

Yushkevich, P.A., Piven, J., Hazlett, H.C., Smith, R.G., Ho, S., Gee, J.C., Gerig, G., 2006. User-guided 3D active contour segmentation of anatomical structures: Significantly improved efficiency and reliability. NeuroImage 31, 1116–1128. 10.1016/j.neuroimage.2006.01.015

Zhao, F., Pan, H., Li, N., Chen, X., Zhang, H., Mao, N., Ren, Y., 2022. High-order brain functional network for electroencephalography-based diagnosis of major depressive disorder. Front. Neurosci. 16, 976229. 10.3389/fnins.2022.976229

