## Supplementary material for "Systematic cross-species comparison of prefrontal cortex functional networks targeted via Transcranial Magnetic Stimulation": SupplementalMaterials - Systematic cross-species comparison of prefrontal cortex functional networks targeted via Transcranial Magnetic Stimulation.pdf

**Supplementary Materials**

*Additional Metrics for Individual Specific TMS Functional Network Analysis*

This study used four metrics to quantify the ability of the TMS-induced electric field to target one of the seven Yeo parcellation networks. The primary metric used within this study is the percentage of overlap between the thresholded binarized functional connectivity maps to the individual functional network parcellations. We calculated overlap as the number of nodes in the maps that overlaid with each functional network divided by the total number of nodes within the functional network. A secondary metric we used to investigate TMS-network activation is the DICE Coefficient. The DICE Coefficient quantified the similarity between the binarized thresholded functional connectivity map and each Yeo parcellation. We also calculated the spatial correlation between the simulated map and the Yeo parcellation network. Finally, we looked at the average functional correlation that overlaps with each Yeo parcellation.

### Supplemental Figures

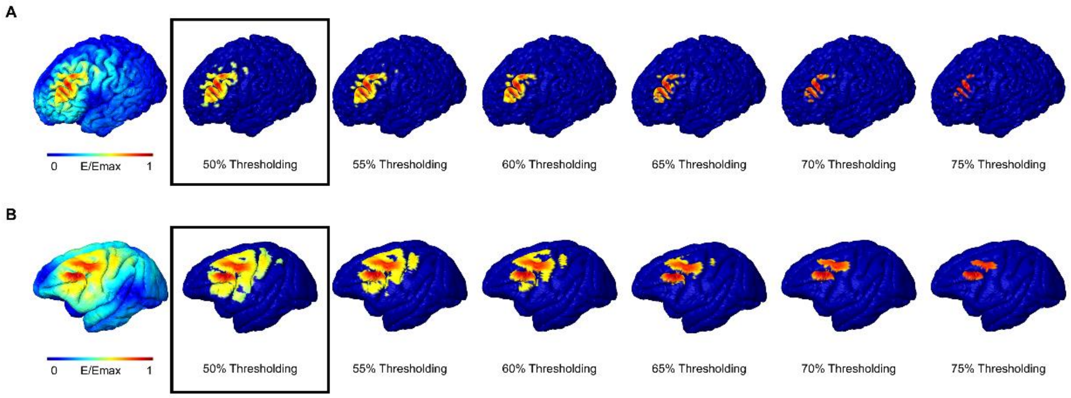


**Supplementary Figure 1. Effect of electric field thresholding. A)** Electric field distribution for one coil location and orientation on human 32k fsLR cortical surface. A threshold is calculated by identifying the robust maximum of the electric field (left) and applying a percentage (50-75%, in 5% increments, shown left to right). All values above this threshold (right) are identified as the seed region for generating the functional connectivity maps. Increased electric field threshold increases the specificity of the seed region. **B)** The electric field distribution on an NHP 32k fsLR cortical surface (left) is shown for one coil configuration (one unique coil location and orientation). The thresholding technique for the NHPs is the same approach outlined for the human models. Like the human models, increasing the electric field thresholding value increases the specificity of the seed region (right).

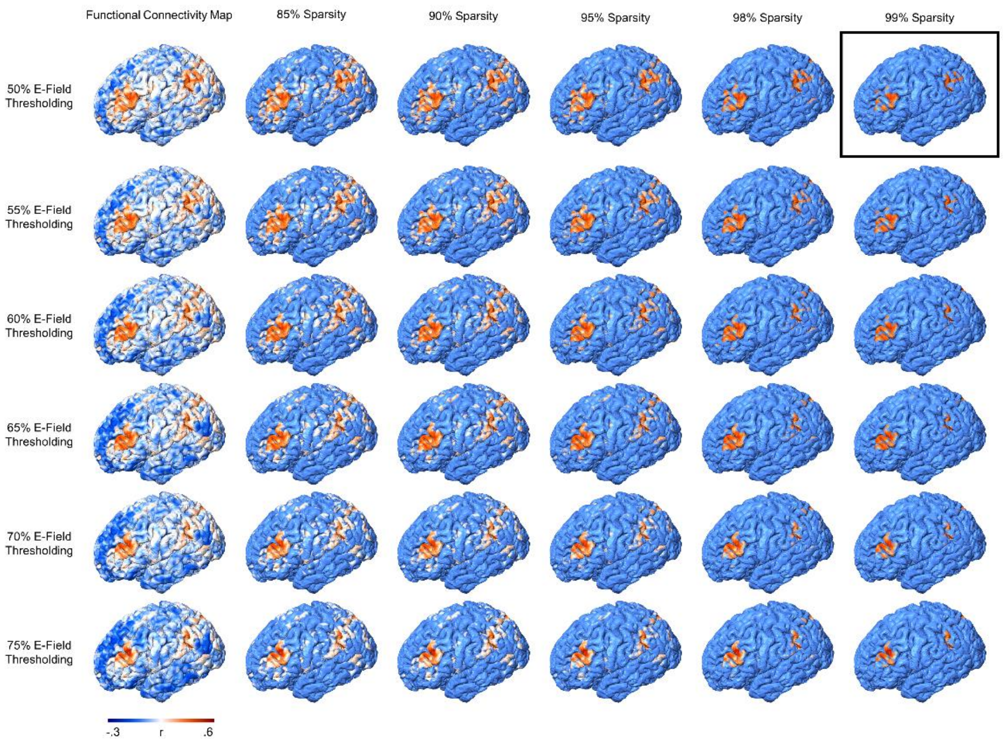


**Supplementary Figure 2. Effect of functional connectivity thresholding on a human cortical surface.** For higher sparsity thresholds, the functional connectivity seed region becomes more focal. The activation hotspots (identified in red) remain spatially the same across all functional and electric field thresholding levels. Increasing the functional connectivity threshold reduces the presence of low-level correlation values in the final network activation analysis. The black box indicates the configuration used within this study.

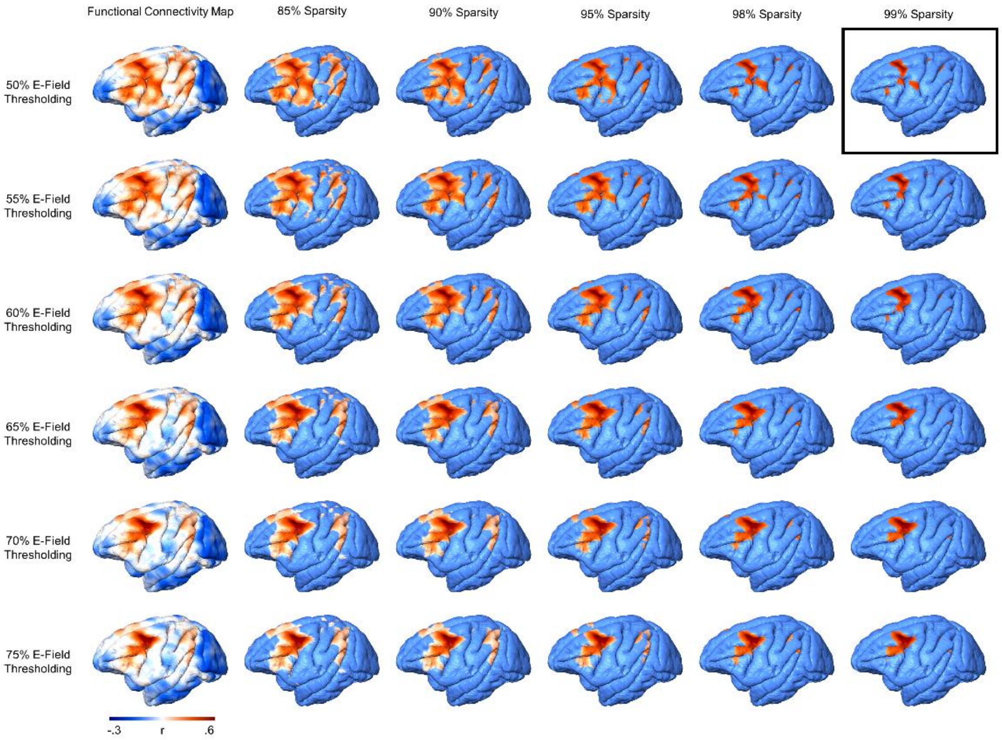


**Supplementary Figure 3. Effect of functional connectivity thresholding on a macaque cortical surface.** For higher sparsity thresholds, the functional connectivity seed region becomes more focal. For all investigated functional and electric field thresholding levels, the activation hotspots (identified in red) remain at the same spatial location. The black box indicates the configuration used within this study.

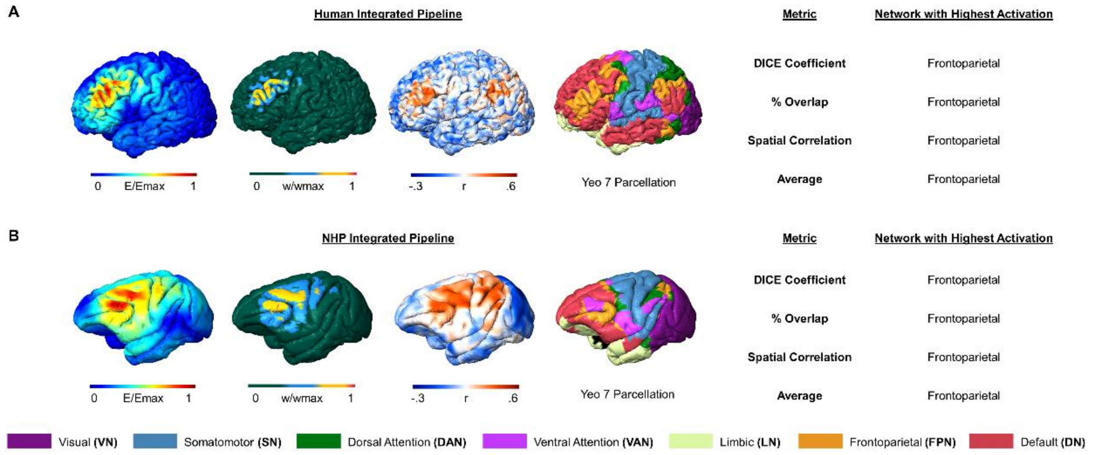


**Supplementary Figure 4. Alternative metrics to identify TMS-targeted functional connectivity networks. A)** Example human integrated TMS-FC targeting pipeline. Four alternative evaluation metrics were used to identify the FC network with the highest network activation (1 – DICE Coefficient, 2 – Percentage of Overlap, 3 – Spatial Correlation, and 4 – Average Activation). All methods identified the Frontoparietal Network as the network with the highest activation for this specific coil location and orientation. **B)** Example NHP integrated TMS-FC targeting pipeline. Four alternative evaluation metrics were used to identify the FC network with the highest network activation (1 – DICE Coefficient, 2 – Percentage of Overlap, 3 – Spatial Correlation, and 4 – Average Activation). All methods identified the Frontoparietal Network as the network with the highest activation for this specific coil location and orientation.

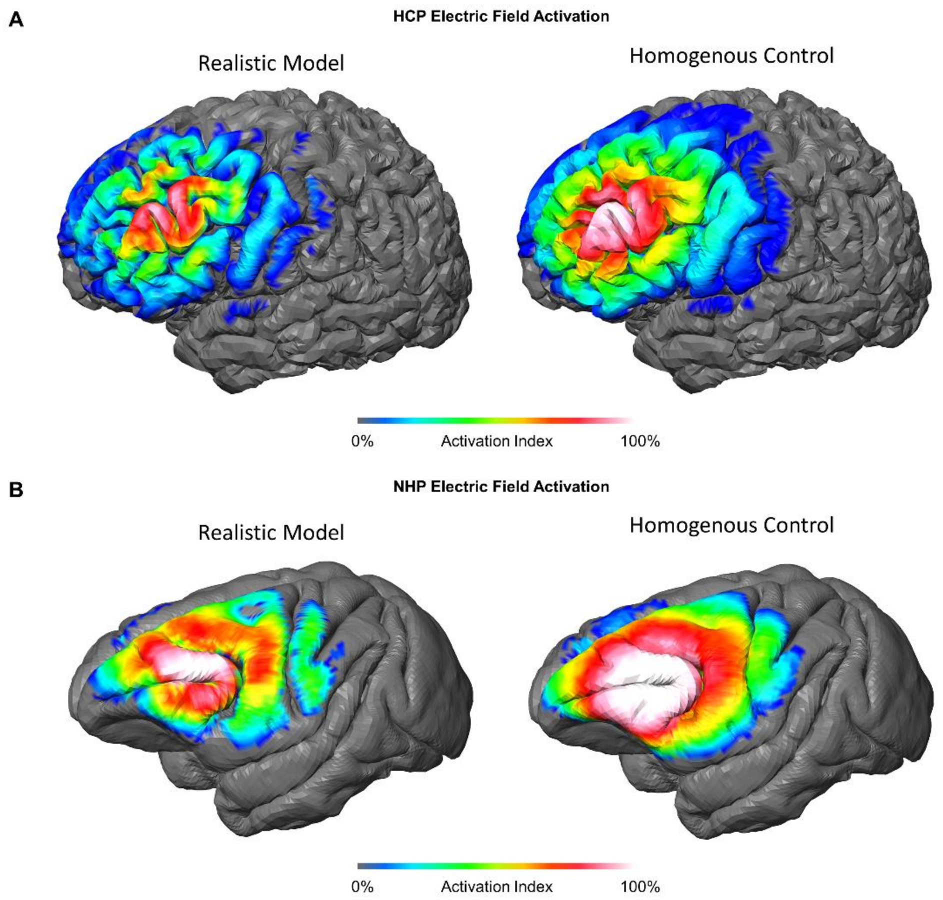


**Supplementary Figure 5. Index of activation for both realistic and homogenous conductivities.** For the realistic conductivity FEM models (left column), the assigned tissue conductivities were $\sigma_{skin} = 0.465S/m$, $\sigma_{skull} = 0.010S/m$, $\sigma_{CSF} = 1.654S/m$, $\sigma_{GM} = 0.276S/m$, $\sigma_{WM} = 0.126S/m$. For the homogenous conductivity model (right column), all tissue conductivities were assigned to $\sigma = 0.276S/m$. **A)** Across all stimulation conditions in the human realistic models, the electric field activation index is distributed across the stimulation grid area with the highest area of activation at gyral crowns in the center grid region. In the homogenous conductivity models, the activation distribution is similar, but with a higher activation index under the center of the grid. **B)** Across all simulation conditions in the NHP realistic models, an activation preference for the defined prefrontal gryi is identified arising from the CSF-GM interface. In the homogenous conductivity model areas of activation are more spread out irrespective of GM anatomy supporting the importance of brain gyrification on targeted brain areas.
